# Organelle intergenic DNA is preferentially distributed between opposite-polarity gene pairs

**DOI:** 10.64898/2026.10.08.757646

**Authors:** Matheus Sanita Lima, Breno C. Dupin, David R. Smith

**Author notes:** These authors contributed equally (Matheus Sanita Lima, Breno C. Dupin).

## Abstract

Across eukaryotes, mitochondrial and plastid genomes vary enormously in size and noncoding content. This variation is often attributed to diverging mutation rates, DNA repair mechanisms, and the power of genetic drift. Although these processes can help explain how much noncoding DNA an organelle genome may retain, they do not explain how this noncoding DNA is distributed throughout the genome. We analyzed over 2.6 million intergenic regions (IGRs) from ∼37,000 mitochondrial and plastid genomes to test whether gene-pair polarity marks local differences in the distribution of organelle intergenic DNA. Taxonomy-structured mixed models, controlling for flanking-gene length and type, showed that IGRs between opposite-polarity gene pairs were 1.1–3.2-fold longer than those between same-polarity genes. IGRs between convergent (+−) and divergent (−+) gene pairs showed distinct residual-dispersion patterns, indicating that they are not homogeneous in their length distribution. The prevalence of opposite-polarity gene pairs showed no consistent relationship with organelle genome size or intergenic DNA fraction. Ultimately then, gene-pair polarity consistently marks where organelle intergenic DNA is preferentially located.

## INTRODUCTION

Mitochondria and plastids contain their own genomes, relics of their prokaryotic past. These genomes, which maintain various prokaryotic-like attributes (e.g., polycistronic transcription), can differ immensely in size and (non)coding proportion: ∼3 kb-12 Mb and ∼1-99% coding (Smith and Keeling 2015). There are various hypotheses to account for this diversity, including the Mutational Hazard Hypothesis (MHH), which argues that differences in organelle genome size and noncoding content can be explained by variation in mutation rates (Lynch et al. 2006). Although the MHH has strong empirical support coming from model species, organelle mutation data remain sparse for non-model eukaryotes, and findings from some lineages are inconsistent with the MHH predictions (Smith 2016). Organelle DNA repair mechanisms have also been invoked to explain genome size variation across organelles, particularly for land plant mitochondria and green algal plastids (Christensen 2014; Smith 2020) but repair mechanisms are not readily amenable to large-scale comparisons.

Furthermore, these prevailing hypotheses address why noncoding DNA content differ among lineages, but they do not explain how and why noncoding nucleotides are often unevenly distributed between genes. Although local sequence contexts and functional constraints can vary (Morton 2022; Lake et al. 2024), IGRs within the same genome should share a broad population-genetic, mutational, and DNA-repair environment, yet can differ substantially in length. Within-genome comparisons can therefore help identify local genomic features associated with IGR length while reducing confounding by properties that vary among genomes and lineages.

Gene-pair polarity, for instance, could represent a local genomic feature impacting IGR length. Same-polarity gene pairs can exist in sense orientation on a polycistronic transcript; however, this is not true for convergent (+−) or divergent (−+) gene pairs. In other words, same-polarity gene pairs are more likely to be transcriptionally coupled compared to opposite-polarity pairs. Consequently, we used gene-pair polarity as a proxy for transcriptional coupling, while still recognizing that this feature alone cannot capture transcript structure perfectly.

We reasoned that transcriptionally coupled genes may share regulatory architecture, whereas opposite-polarity genes represent transcriptionally distinct boundaries that may be subject to different spatial constraints. Accordingly, we hypothesized that IGRs between same-polarity gene pairs would be shorter than those between convergent or divergent gene pairs. Relationships between gene orientation and intergenic spacing have been explored in bacterial, archaeal, and nuclear genomes (Gilet et al. 2020; Huber et al. 2023), but, to our knowledge, have not been systematically examined across organelle genomes. Here, we show that IGRs between opposite-polarity gene pairs are consistently longer than those between same-polarity pairs across mitochondrial and plastid genomes. Convergent and divergent IGRs exhibit distinct, lineage-dependent length distributions, which may reflect the underlying regulatory architecture. Despite being preferential loci for genomic expansion, the prevalence of opposite-polarity gene pairs shows no consistent relationship with organelle genome size or noncoding content. Together, these results identify gene-pair polarity as an important local correlate of intergenic DNA distribution rather than the key determinant of genome-wide noncoding content.

## RESULTS AND DISCUSSION

### Opposite-polarity gene pairs disproportionately delimit long intergenic regions

We extracted 2,637,725 IGRs from 37,002 mitochondrial and plastid genomes of plants, green algae, protists, fungi, and metazoans. Within genomes, opposite-polarity gene pairs delimited the longer IGRs in ∼85% of genomes overall (**Figure 1A; Table S1**), including 99.9% of plant plastid and 94.3% of plant mitochondrial genomes. In genomes with ≥ 40 IGRs, opposite-polarity gene pairs were enriched among the longest IGRs in 96.6% of genomes and were depleted among the shortest IGRs in 98% (**Table S1**). When pooling all IGRs within each taxonomic group and organelle type, same-polarity IGRs occupied less intergenic sequence than expected from their count alone, whereas divergent IGRs were enriched in all groups (**Figure 1B**). Convergent IGRs were enriched in most groups but were approximately proportional in plant plastids and depleted in metazoan mitochondria.

**Figure 1.**
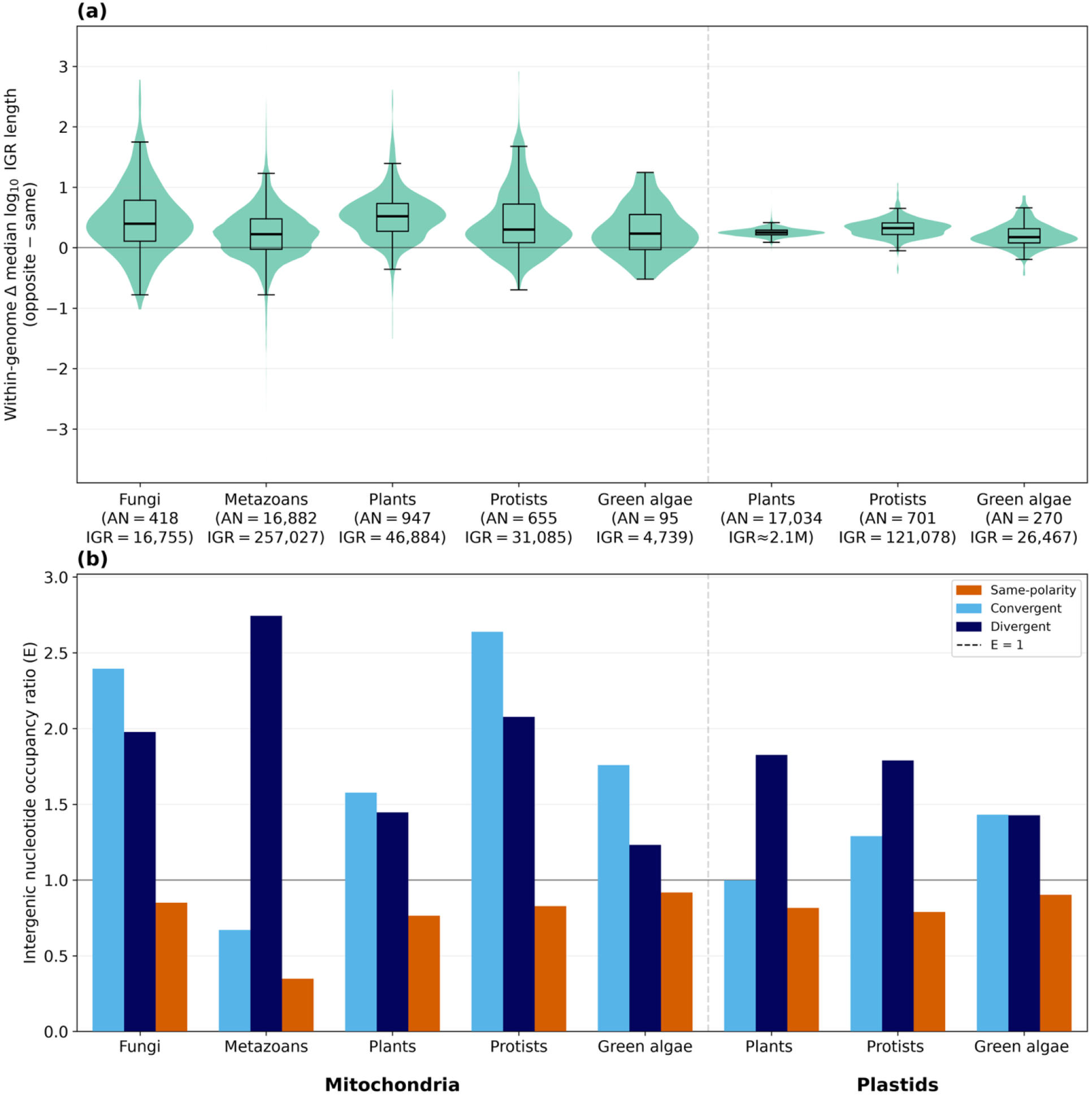
Opposite-polarity gene pairs delimit longer intergenic regions and account for a disproportionate share of intergenic sequence. a) Distribution among genomes of the within-genome difference in median log_10_-transformed intergenic-region (IGR) length between opposite- and same-polarity gene pairs, calculated as Δ = median[log_10_(*L*opposite)] − median[log_10_(*L*same)], where *L* = length. Opposite-polarity IGRs are delimited by convergent (+−) and divergent (−+) gene pairs, whereas same-polarity IGRs are delimited by ++ and −− gene pairs. Violin widths represent the density of genome-level Δ values, and embedded boxplots summarize their median and interquartile range. The horizontal line at Δ = 0 indicates equal median lengths. Numbers below each dataset indicate the respective numbers of analyzed organelle genome sequences and IGRs. (B) Intergenic-nucleotide occupancy ratio (E) for each polarity class, calculated after pooling IGR counts and summed lengths within each dataset, so bars represent pooled descriptive ratios. Vertical dashed lines separate mitochondrial and plastid datasets. Only genomes containing at least one IGR in each of the three polarity classes were included.

The two opposite-polarity classes also differed in their relative lengths. At the pooled descriptive level, median convergent IGR lengths exceeded median divergent IGR lengths in all five mitochondrial datasets by 1.06-to 1.91-fold, whereas median divergent IGR lengths exceeded median convergent IGR lengths in all three plastid datasets by 1.38-to 2.82-fold (**Table S1**). Thus, although convergent and divergent gene pairs both tended to delimit longer IGRs than same-polarity pairs, their relative length distributions differed between mitochondrial and plastid genomes. Opposite-polarity gene pairs therefore delimit a disproportionate share of intergenic sequence wherever they occur; an association observed across organelle genomes spanning roughly three orders of magnitude in size. Together, these descriptive analyses established a consistent (unadjusted) association between gene-pair polarity and IGR length. Because the lengths and types of the flanking genes could differ among polarity classes and could themselves be associated with IGR length, we next asked whether these features accounted for the observed association.

### The association between gene-pair polarity and IGR length persists after accounting for flanking-gene architecture

We fitted taxonomy-structured Bayesian linear mixed models to estimate the association between gene-pair polarity and IGR length while accounting for flanking-gene architecture. The models included log_10_ IGR length as the response (modeled with a Gaussian likelihood and identity link); gene-pair polarity as the effect of interest; maximum flanking-gene length and flanking-gene type as covariates; and a taxon-level random intercept structured by a taxonomy-derived covariance matrix. Both dimensions of flanking-gene architecture contributed complementary predictive information: in all eight groups, the full model had higher expected log predictive density under three-fold cross-validation than models containing either flanking-gene length or flanking-gene type alone (**Table S2**). Nevertheless, the estimated polarity-associated fold differences from the full models remained similar to those from the polarity-only models, changing by only ∼ −19% to +6%. Across all eight groups, convergent and divergent IGRs were estimated to be 1.1-to 3.2-fold longer than same-polarity IGRs, with all 95% credible intervals excluding 1 (**Figure 2**). The adjusted differences ranged from a 1.08-fold increase for convergent IGRs in protist plastids to a 3.17-fold increase for divergent IGRs in the same group. Thus, although flanking-gene length and type contributed to predictive performance, neither accounted for the association between gene-pair polarity and IGR length.

**Figure 2.**
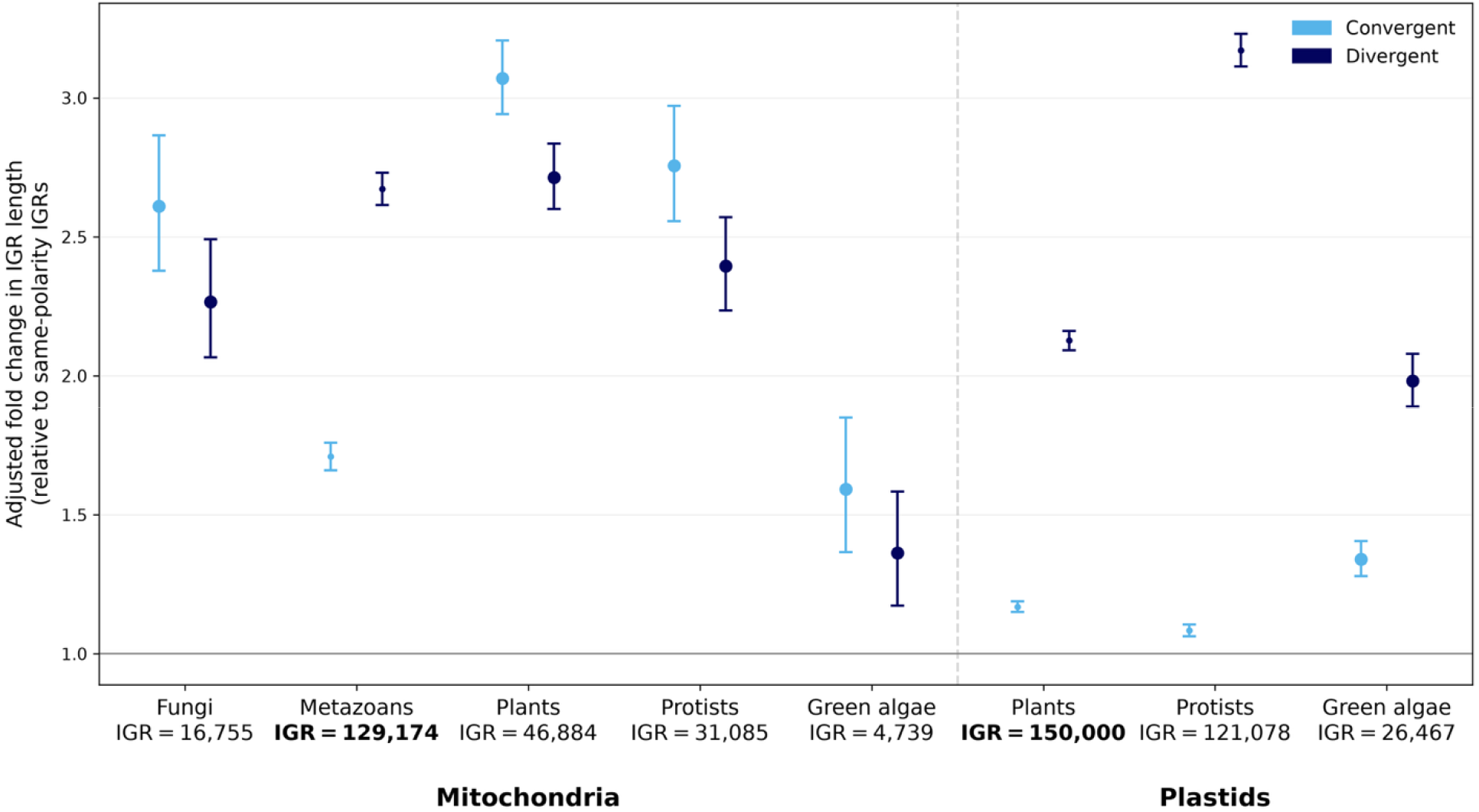
The association between gene-pair polarity and intergenic region length persists after accounting for flanking-gene architecture. Posterior estimates from taxonomy-structured Bayesian linear mixed models fitted separately to the eight datasets. Models included maximum flanking-gene length and flanking-gene type as covariates. Points show posterior median adjusted intergenic-region (IGR) length ratios for convergent (+−) and divergent (−+) IGRs relative to same-polarity (++ and −−) IGRs. Error bars show central 95% credible intervals. Ratios were obtained by exponentiating the polarity coefficients as 10^β^. The horizontal line at 1 indicates no adjusted difference from same-polarity IGRs. Numbers below each dataset indicate the number of IGRs included in the corresponding model. Bolded numbers indicate datasets that were subjected to genus-level filtering and down sampling. The vertical dashed line separates mitochondrial and plastid datasets.

### Convergent and divergent IGRs differ in relative length and dispersion

As convergent and divergent gene pairs represent different transcriptional configurations, we asked whether they also differed in residual IGR-length variability. We fitted distributional models in which residual dispersion was allowed to vary among polarity classes. Overall, convergent IGRs were more variable than same-polarity IGRs in six groups (σ-ratio up to 1.29), whereas divergent IGRs showed similar or lower dispersion in seven groups (σ-ratio as low as 0.65; **Table S3**). These asymmetric patterns indicate that convergent and divergent IGRs do not form a homogeneous opposite-polarity class. Notably, residual dispersion is not a direct measure of evolutionary constraint. Differences in residual variability could instead arise from unmeasured, lineage-specific variation in regulatory architecture or mutational processes, including repeat-associated insertions and deletions (Wagle et al. 2001; Kim and Kim 2014), among other mechanisms. The dispersion contrasts therefore establish distinct, lineage-dependent length distributions, while the mechanisms generating those distributions remain unresolved.

Direct posterior contrasts of the adjusted effects showed that divergent IGRs were longer than convergent IGRs in plastid genomes, with divergent-to-convergent length ratios ranging from 1.5 to 2.9 (**Table S4**). Mitochondrial genomes were more heterogeneous: convergent IGRs were longer in fungi, plants, and protists. Green-algal mitochondria showed a similar point estimate, although the 95% credible interval included no statistical difference (divergent-to-convergent ratio: 0.9, 0.7–1.0). Metazoan mitochondria showed the reverse pattern, with divergent IGRs estimated to be 1.6-fold longer than convergent IGRs (1.5–1.6).

Regulatory and transcriptional architecture may contribute to these differences. In plastid systems, promoter-proximal sequences influence transcription initiation, whereas 5′-UTR sequences affect transcript accumulation and translation (Eibl et al. 1999; Castandet et al. 2019; Puthiyaveetil et al. 2021). Where transcription termination is inefficient, mature transcript 3′ ends are formed and stabilized post-transcriptionally through RNA processing and the activity of RNA-binding proteins (Hayes et al. 1996; Rott et al. 1996; Sharwood et al. 2011; Castandet et al. 2019). The spatial requirements of these regulatory and processing features might contribute to the consistent divergent-length excess observed in plastids, although their effects cannot be assumed to be uniform across the diverse plastid lineages analyzed here. In fact, heterogeneity in both adjusted length and residual-dispersion contrasts among datasets suggests that a uniform contribution from fixed regulatory elements is unlikely to provide a complete explanation. Variation in polycistronic transcription and intercistronic RNA processing can alter transcript organization and expression (Zhou et al. 2007; Guilcher et al. 2021), whereas repeat-rich intergenic regions can expand through repeat-associated indels and sequence rearrangements (Wagle et al. 2001; Kim and Kim 2014). These and other lineage-specific aspects of genome architecture likely contribute to the observed length distributions.

### Gene-pair polarity marks local distribution of intergenic DNA, not genome-wide abundance

Having established a local association between gene-pair polarity and IGR length, we tested whether the prevalence of opposite-polarity gene pairs also tracked genome size or noncoding fraction. Across datasets, the per-genome prevalence of opposite-polarity gene pairs showed no consistent relationship with genome size (Spearman’s ρ from −0.19 in green-algal plastids to 0.68 in protist plastids) or noncoding content (−0.08 to 0.36; **Figure 3; Table S5**). Gene-pair polarity, therefore, marks the local distribution of intergenic DNA without consistently explaining genome-wide variation in noncoding content. These findings complement the prevailing models of genome evolution in which mutation, genetic drift, and DNA repair mechanisms explain how much noncoding DNA a genome retains (Lynch et al. 2006; Christensen 2014; Smith 2020).

**Figure 3.**
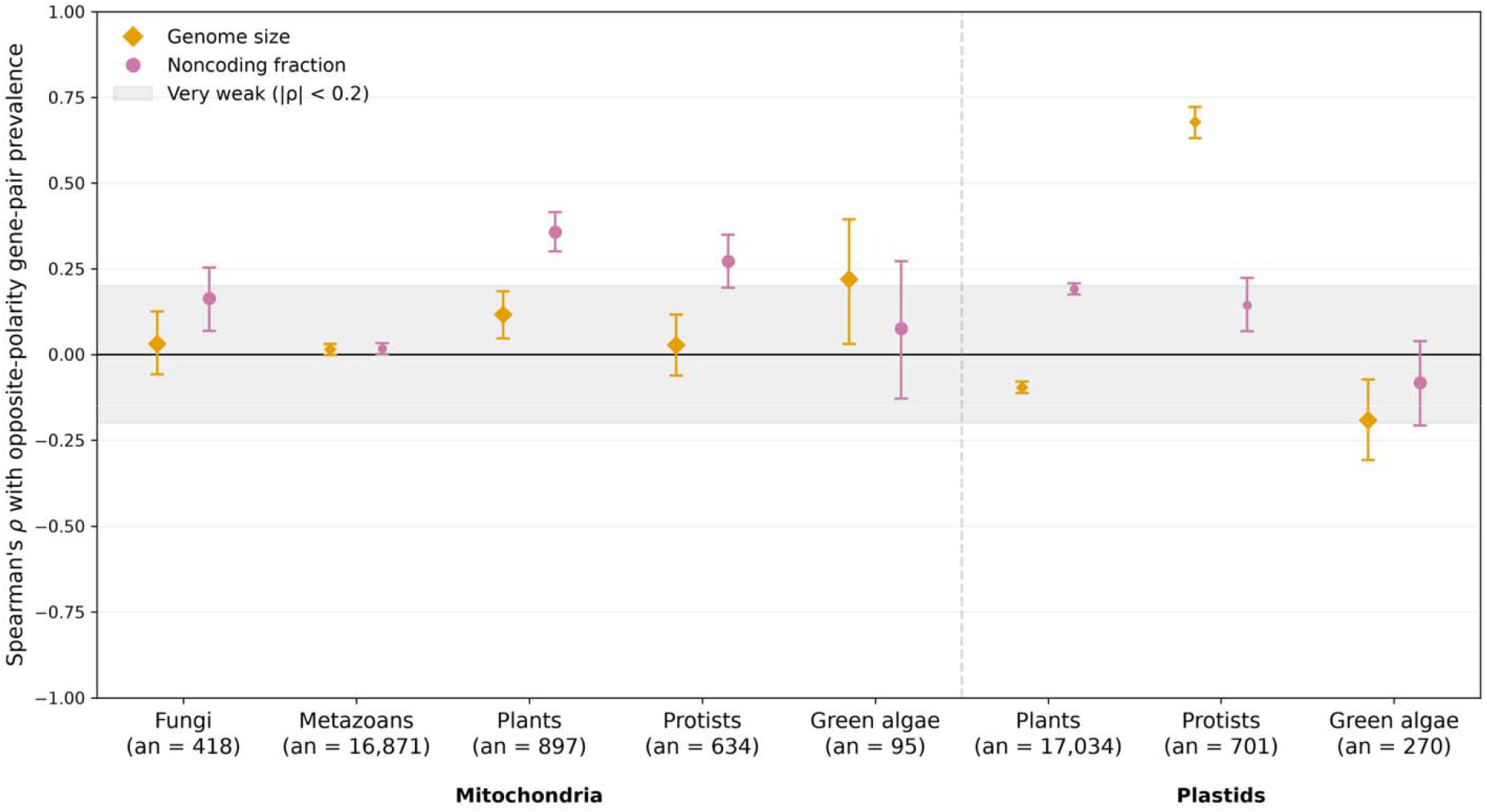
Opposite-polarity gene-pair prevalence shows no consistent relationship with genome size or intergenic noncoding fraction. Points show Spearman’s rank correlation coefficient (ρ) between the per-genome prevalence of opposite-polarity gene pairs and either genome size or intergenic noncoding fraction within each dataset. Opposite-polarity prevalence was calculated as the number of convergent (+−) and divergent (−+) IGRs combined, divided by the total number of IGRs in each genome. Intergenic noncoding fraction was calculated as the summed length of all extracted IGRs divided by total genome length. Error bars show percentile 95% bootstrap confidence intervals from 2,000 replicates. The horizontal line at ρ=0 indicates no monotonic association, and the shaded band marks very weak correlations (∣ρ∣<0.2). Numbers below each dataset indicate the number of organelle genome sequences included in the analysis. The vertical dashed line separates mitochondrial and plastid datasets. Analyses were restricted to genomes containing at least five IGRs.

Our analyses do not establish that changes in gene orientation necessarily cause IGRs to expand. Alternative promoters, transcript processing, and pervasive read-through transcription can also decouple some same-polarity genes, making polarity an informative but imperfect proxy for transcriptional coupling (Kühn et al. 2005; Zhou et al. 2007; Sharwood et al. 2011; Sanita Lima and Smith 2017). Comparative analyses of lineages with independently derived gene-order rearrangements, together with long-read transcriptomic studies linking transcript continuity/coverage to IGR length, could test whether the observed patterns arise directly from transcriptional architecture or from other features associated with gene orientation. Nevertheless, the recurrence of the association across structurally diverse mitochondrial and plastid genomes, encompassing markedly different transcriptional architectures (Smith and Keeling 2016; Shtolz and Mishmar 2023), highlights a recurring distinction between the local distribution and genome-wide abundance of intergenic nucleotides. Opposite-polarity gene pairs consistently delimit longer IGRs, while convergent and divergent IGRs exhibit distinct, lineage-dependent mean and dispersion patterns. Gene-pair polarity marks the local distribution of intergenic DNA without consistently explaining genome-wide variation in intergenic content across lineages.

## METHODS

### Genome acquisition and quality control

Mitochondrial and plastid genome records were obtained from NCBI Nucleotide (Sayers et al. 2025) and NCBI Organelle (O’Leary et al. 2024), with GFF3 annotations and FASTA sequences retrieved using custom Python scripts. Genomes were divided into eight datasets according to organelle type and NCBI taxonomic classification (Schoch et al. 2020): fungal, metazoan, plant, protist, and green-algal mitochondria, and plant, protist, and green-algal plastids. Records yielding empty or deprecated files were removed, as were genomes whose GFF3 files lacked both tRNA and rRNA annotations. Complete lists of accession numbers, including those excluded during quality control, are provided in the project repository.

### Intergenic region extraction and classification

After quality control, intergenic regions (IGRs) were extracted using GDT (Gene Dictionary Tool) 1.0.3 and TIGRE 2.0.3 following published protocols (Dupin et al. 2025a, 2025b). Each IGR was classified according to the polarity of its two flanking genes as same-polarity (++ or −−), convergent (+−), or divergent (−+). Convergent and divergent IGRs were collectively classified as opposite-polarity for descriptive analyses. Only genomes containing at least one IGR in each of the three polarity classes were retained for downstream analysis. IGRs located within plastid inverted repeats were not deduplicated.

### Descriptive analyses

For each genome, we calculated the within-genome difference between the median log_10_-transformed lengths of opposite- and same-polarity IGRs: Δ=median[log10(*L*opposite)]-median[log10(*L*same)]. Length-tail enrichment was computed for genomes with ≥40 IGRs. For each tail, we divided the fraction of opposite-polarity IGRs among the longest or shortest 10% of a genome’s IGRs by the genome-wide opposite-polarity fraction. To quantify the amount of intergenic sequence occupied by each polarity class, IGR counts and summed lengths were pooled within each dataset. The occupancy ratio was calculated as E = P_obs / P_exp, where P_obs is a class’s proportion of total intergenic length and P_exp is its proportion of the total IGR count. Thus, E>1 indicates that a polarity class occupies more intergenic sequence than expected from its numerical frequency.

### Taxonomy-structured mixed models

For each dataset, we fitted Bayesian linear mixed models using brms version 2.23.0 (Bürkner 2017), which interfaces with Stan for Bayesian inference (Carpenter et al. 2017). Log_10_-transformed IGR length was modeled using a Gaussian likelihood and identity link. Four mean structures were considered: (i) gene-pair polarity only; (ii) polarity and maximum flanking-gene length; (iii) polarity and flanking-gene type; and (iv) polarity and both flanking-gene covariates. Maximum flanking-gene length was log_10_-transformed and centered on its mean in the dataset used for model fitting. Protein-coding gene–gene pairs were the reference flanking-gene type, and same-polarity IGRs were the reference polarity class. For each dataset, a taxonomic topology containing the represented NCBI taxon identifiers was obtained using ETE 4 (Huerta-Cepas et al. 2016) and an NCBI Taxonomy database (updated on July 7, 2026). The topology was converted to an ultrametric tree using `to_ultrametric()`. In R, polytomies were resolved as zero-length dichotomies using ape::multi2di, and the tree was pruned to the taxa retained in the analyzed dataset. Matrix (A) was calculated as the correlation form of the among-taxon covariance matrix using ape::vcv(tree, corr = TRUE) (Paradis and Schliep 2019). All models included a taxon-level random intercept structured by matrix (A), specified as `(1 | gr(ncbi_taxid, cov = A))`.

Before model fitting, the “metazoan mitochondria” and “plants plastid” datasets were restricted to the first genome record per genus, with genus assigned from the NCBI organism name. The plant plastid dataset was then reduced to 150,000 IGRs by proportional stratified random sampling within polarity classes, using a random seed of 42. Models were fitted using four chains of 6,000 iterations each, including 2,000 warm-up iterations, yielding 16,000 post-warm-up draws. We used `adapt_delta = 0.95`, `max_treedepth = 12`, and the default priors generated by `brms`. Effects of convergent and divergent polarity relative to the same-polarity reference were exponentiated as 10^β^ and are reported as adjusted length ratios with 95% credible intervals.

### Model comparison, dispersion, and direct contrasts

The three covariate-containing mean structures were compared using expected log predictive density under three-fold cross-validation (Vehtari et al. 2017), implemented with `kfold` and `loo_compare`. Within each NCBI taxon, IGRs were randomly distributed approximately evenly among the three folds. Each cross-validation refit used one chain of 6,000 iterations, including 2,000 warm-up iterations. To test whether residual IGR-length variability differed among polarity classes, we fitted a distributional model with the both-covariate mean structure and `sigma ∼ polarity_3bin`. Because sigma was modeled using a log link, exponentiated sigma coefficients represent residual-standard-deviation ratios relative to same-polarity IGRs on the log_10_-length scale. Across all full-data mean and distributional fits, rank-normalized split-R-hat was ≤1.01 for every parameter, with no divergent transitions and no iterations reaching the maximum tree depth (Vehtari et al. 2021). For each covariate-containing mean model, the proportion of residual variance attributable to the taxonomy-structured random intercept was summarized from posterior draws as ICC_taxon=σ_taxon^2/(σ_taxon^2+σ_residual^2). Convergent and divergent IGRs were compared directly using their joint posterior draws extracted and summarized with the posterior package version 1.6.1 (Bürkner et al. 2025). For each draw, the adjusted mean contrast was calculated as Δmean = βdivergent−βconvergent, and 10^Δmean^ gives the adjusted divergent-to-convergent length ratio. Contrasts were summarized using posterior medians, central 95% credible intervals, and directional posterior probabilities.

### Prevalence associations

For each genome, opposite-polarity prevalence was calculated as the number of convergent and divergent IGRs divided by the total number of IGRs. Genome size was obtained from the corresponding genome record, and intergenic fraction was calculated as the summed length of all extracted IGRs divided by genome length. Analyses were restricted to genomes containing at least five IGRs. Within each dataset, Spearman’s ρ was calculated between opposite-polarity prevalence and (i) genome size and (ii) intergenic fraction. Kendall’s τ-b was calculated as a rank-based robustness measure. Percentile 95% confidence intervals for both coefficients were obtained from 2,000 bootstrap replicates in which genomes were sampled with replacement and paired variables were resampled jointly. Nonfinite paired observations were excluded before analysis, and a random seed of 42 was used. Because the bootstrap did not explicitly account for taxonomic non-independence, the correlations were interpreted as descriptive genome-level associations.

## AUTHOR CONTRIBUTIONS

Conceptualization, M.S.L. and D.R.S.; methodology, B.C.D. and M.S.L.; Investigation, M.S.L and B.C.D.; writing – original draft, M.S.L. and B.C.D.; writing – review & editing, M.S.L., B.C.D. and D.R.S.; funding acquisition, D.R.S.; resources, D.R.S.; supervision, D.R.S.

## DATA AVAILABILITY

The data and code underlying this article are available in Zenodo, at https://zenodo.org/records/21540121. The project repository is https://gitlab.com/brenodupin/datasets-organelle-igr.

## ACKNOWLEDGMENTS

The authors are thankful to Kevin Fontyn for statistical consultation. D.R.S. is supported by the Natural Sciences and Engineering Research Council of Canada (NSERC). The authors have used Artificial Intelligence tools (ChatGPT and Claude) for: i) code optimization and refactoring; and ii) manuscript copyediting and proof-reading.

## SUPPLEMENTARY MATERIAL

Supplementary material is available at Molecular Biology and Evolution online.

